# Control of aperture closure during reach-to-grasp movements in immersive haptic-free virtual reality

**DOI:** 10.1101/2020.08.01.232470

**Authors:** Madhur Mangalam, Mathew Yarossi, Mariusz P. Furmanek, Eugene Tunik

## Abstract

Virtual reality (VR) has garnered much interest as a training environment for motor skill acquisition, including for neurological rehabilitation of upper extremities. While the focus has been on gross upper limb motion, VR applications that involve reaching for, and interacting with, virtual objects are growing. The absence of true haptics in VR when it comes to hand-object interactions raises a fundamentally important question: can haptic-free immersive virtual environments (hf-VEs) support naturalistic coordination of reach-to-grasp movements? This issue has been grossly understudied, and yet is of significant importance in the development and application of VR across a number of sectors. In a previous study (Furmanek et al. 2019), we reported that reach-to-grasp movements are similarly coordinated in both the physical environment (PE) and hf-VE. The most noteworthy difference was that the closure phase—which begins at maximum aperture and lasts through the end of the movement—was longer in hf-VE than in PE, suggesting that different control laws might govern the initiation of closure between the two environments. To do so, we reanalyzed data from Furmanek et al. (2019), in which the participants reached to grasp three differently sized physical objects, and matching 3D virtual object renderings, placed at three different locations. Our analysis revealed two key findings pertaining to the initiation of closure in PE and hf-VE. First, the respective control laws governing the initiation of aperture closure in PE and hf-VE both included state estimates of transport velocity and acceleration, supporting a general unified control policy for implementing reach-to-grasp across physical and virtual environments. Second, aperture was less informative to the control law in hf-VE. We suggest that the latter was likely because transport velocity at closure onset and aperture at closure onset were less independent in hf-VE than in PE, ultimately resulting in aperture at closure onset having a weaker influence on the initiation of closure. In this way, the excess time and muscular effort needed to actively bring the fingers to a stop at the interface of a virtual object was factored into the control law governing the initiation of closure in hf-VE. Critically, this control law remained applicable, albeit with different weights in hf-VE, despite the absence of terminal haptic feedback and potential perceptual differences.

## Introduction

Virtual reality (VR) has garnered much interest as a training environment for motor skill acquisition, including for neurological rehabilitation of upper extremities (Schultheis and Rizzo 2001; Sveistrup 2004; Holden 2005; Rizzo and Kim 2005; Rose et al. 2005; Adamovich et al. 2009; Cheung et al. 2014). The ability to reach-to and grasp virtual objects is paramount to any VR-based manual training. Devices that provide haptic feedback in VR tend to be expensive, bulky, and restrict natural motion in the virtual environment (Borst and Volz 2005; Pacchierotti et al. 2017; Culbertson et al. 2018). Haptic-free immersive virtual environments (hf-VEs) are a more viable option. However, the extent to which hf-VEs support naturalistic coordination of reach-to-grasp movements is mostly unknown. To fill this gap, in a previous study (Furmanek et al. 2019), we asked participants to reach to grasp physical objects of different sizes located at different distances, and to grasp virtual renderings of these objects, in an immersive hf-VE. We found that reach-to-grasp movements are similarly coordinated in both the physical environment (PE) and immersive hf-VE. The most noteworthy difference was that the closure—which begins at maximum aperture and lasts through the end of the movement—was longer in hf-VE than in PE. Understanding the factors underlying this difference is essential to modify a given hf-VE so that it provides for more naturalistic reach-to-grasp movements.

Much attention has been given to the processes that govern/trigger the initiation of closure during reach-to-grasp movements. Based on the observation that maximum aperture occurs approximately when hand transport deceleration is maximum, it was initially suggested that the initiation of closure is triggered by the reduction of hand transport velocity (Jeannerod 1984; Paulignan et al. 1991a, b). This suggestion was expanded by Rand and colleagues (2006a, b, 2008), who proposed that the closure is initiated based on the dynamics of the arm and hand in relation to the object location and size. According to this view, the distance of the hand from the object at closure is predicted by a linear function combining the state estimates of transport velocity, transport acceleration, and aperture. This control law has been shown to adequately describe the initiation of closure across different movement speeds (Rand et al. 2006b), object distances (Rand et al. 2010), and disease states, such as Parkinson’s disease (Rand et al. 2006a).

To our knowledge, a comparison of the control laws governing the initiation of closure in PE and hf-VEs has not been made before despite reasons to believe that altered perceptual information and/or task constraints in hf-VEs might affect the planning, execution, and control of manual actions (Harris et al. 2019). For example, egocentric depth cues in VE may be processed differently by brain circuits than the natural depth cues in the real world, contributing to uncertainty in perceptual estimates of the object size and distance. Specific to hf-VEs, an absence of terminal haptic feedback may increase uncertainty in perception of the object size as haptics cannot be used to calibrate grasp (Bingham et al. 2007; Fukui and Inui 2013; Renner et al. 2013). Furthermore, the control law governing the initiation of closure in hf-VE might also differ because in PE, the closure is stopped instantaneously as soon as the fingers contact a physical object, but when reaching to grasp a virtual object, voluntary effort must be exerted to arrest finger movement to prevent interpenetration into that object (Prachyabrued and Borst 2012). Knowledge about whether lack of, or uncertainty about, key perceptual information in hf-VE compared to PE changes the control law used to govern closure is crucial as we aim to improve the fidelity of VR applications for manual skills training and rehabilitation.

In the present study, we reanalyzed data from Furmanek et al. (2019) to investigate whether the same control law governs the initiation of closure for reach-to-grasp movements performed in PE and our immersive hf-VE. We hypothesized that slower closure initiated farther from the object in the hf-VE than the PE condition that was noted by Furmanek et al. (2019) resulted from a difference in the control law governing the initiation of closure between the two environments.

To investigate the source of variation in the control low governing the initiation of closure in PE and hf-VE, we followed an information-theoretic approach to model selection (Burnham and Anderson 2002). This approach chooses the best model from a given set of a priori candidate models—by comparing the models against each other, instead of the null hypothesis, and balances the goodness of fit with simplicity. We considered the null model (containing only the intercept) and all the different combinations of the state estimates (aperture, transport velocity, and transport acceleration) to test whether the control law relies on each of these estimates in both PE and hf-VE and whether the strength of reliance on each of these estimates differs between the two environments.

## Methods

### Participants

Thirteen right-handed subjects (2 women; mean±1 *SD* age: 23.9±6.8 years old) with no reported muscular, orthopedic, or neurological health concerns, voluntarily participated in Furmanek et al. (2019) after providing verbal and written consent. Data from these participants were reanalyzed. The experimental task and procedure are described in brief below. Further details about the experimental task and procedure can be found in Furmanek et al. (2019).

### Experimental task

The participants performed the task of reaching toward and grasping three differently-sized physical objects and matching 3D virtual renderings. The objects were cuboids of identical height (8 cm) and depth (2.5 cm) but different widths: small: 3.6 cm; medium: 5.4 cm; large: 7.2 cm (Fig. 1). The physical objects were 3D printed using PLA thermoplastic (mass: small: 30 g; medium: 44 g; large: 59 g) and covered with glow-in-the-dark paint. The participants viewed blue-colored virtual renderings of these objects in a custom 3D immersive hf-VE designed in UNITY (ver. 5.6.1f1, Unity Technologies, San Francisco, CA) and presented via head-mounted display (Oculus Rift DK2, Oculus Inc., Menlo Park, CA). The objects, in PE or hf-VE, were placed at three different distances from the starting position—24 cm (near), 30 cm (middle), 36 cm (far)—and oriented at a 65° angle along the vertical axis to minimize excessive wrist extension (Fig. 1).

**Fig. 1.**
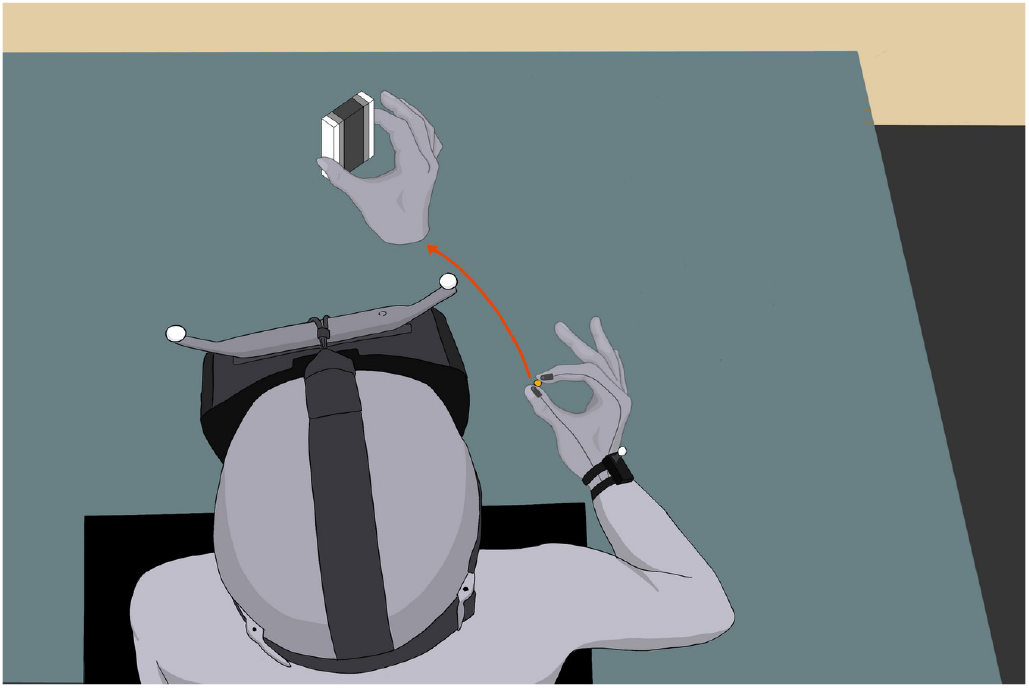
Schematic illustration of the experimental setup and procedure. After wearing an Oculus^TM^ head-mounted display (HMD), the participants sat on a chair in front of the experimental rig, with their thumb pressing a switch (indicated in yellow). IREDs markers were attached to the participant’s wrist, and the tips of the thumb and index finger. An auditory cue —a beep—signaled the participant to reach to grasp the object: black (small: 3.6×2.5×8 cm), dark gray (medium: 5.4×2.5×8 cm), and light gray (large: 7.2×2.5×8 cm), placed at three different distances relative to the switch (near: 24 cm; middle: 30 cm; and far: 36 cm).

### Experimental procedure

The participants were seated in a chair with the right arm and hand placed on a table in front of them (Fig. 1). At the start position, the thumb and index finger straddled a 1.5 cm wide wooden peg located 12 cm in front and 24 cm to the right of the sternum, with the thumb depressing a switch. Lifting the thumb off the switch marked movement onset. A six-camera motion tracking system (PPT Studio N^TM^, WorldViz, Inc., Santa Barbara, CA; sampling rate: 75 Hz) recorded the 3D motion of IRED markers attached to the participant’s wrist (at the center of the segment running between the ulnar and radial styloid process), thumb nail, and index fingernail. A pair of IRED markers attached to the HMD co-registered the participant’s head motion to render the virtual environment. When reaching to grasp a virtual object in hf-VE, the participants viewed the tips of their thumb and index finger as two 3D spheres (green in color, 0.8 cm diameter), reflecting the real-time 3D position of the respective IRED marker. When reaching to grasp a physical object, all lights were turned off so that only the illuminated IRED markers attached to the thumb and index finger, as well as the glow-in-the-dark object, were visible to the participants. The overhead light was turned on after every block of 12 trials to prevent acclimatization of the vision to the dark environment.

Prior to the experiment, the participants practiced the reach-to-grasp task in hf-VE across 135 trials (3 sizes × 3 distances × 15 trials). The experiment consisted of 216 trials, each trial lasting 3 seconds: 108 trials in PE and 108 trials in hf-VE (3 sizes × 3 distances × 12 trials in each environment). The trials were blocked for each size-distance combination and each environment. The order of the presentation of the three objects and their placement was randomized for each block of size-distance combination.

In each trial, an auditory cue—a beep—signaled the participants to reach for, grasp, and lift the object. The participants were considered to have grasped a physical object the moment the tips of their thumb and index finger came into contact with that object’s lateral surface, and aperture did not change more than 1 mm for at least 50 ms. The participants were considered to have grasped a virtual object the instance a custom collision detection algorithm detected that both 3D spheres had come in contact with the virtual object. Once grasp of the virtual object was detected, the object changed color from blue to red. The participants lifted and raised each object briefly before returning their hand to the starting position. The participants were not required to wait for the object to change color before lifting it. The object color change in VE occurred instantaneously when the collision criteria were met. However, because it was impossible to lift the object until the collision criteria were met, it was impossible to lift the object before its color changed. Participants were explicitly instructed to lift the object straight up to discourage forward velocity at the time of grasp. As is shown in Fig. 2C, the planar transport velocity was near zero at the time of grasp in both PE and hf-VE, indicating that the hand transport was halted at the time of grasp, before the upward-lifting motion.

**Fig. 2.**
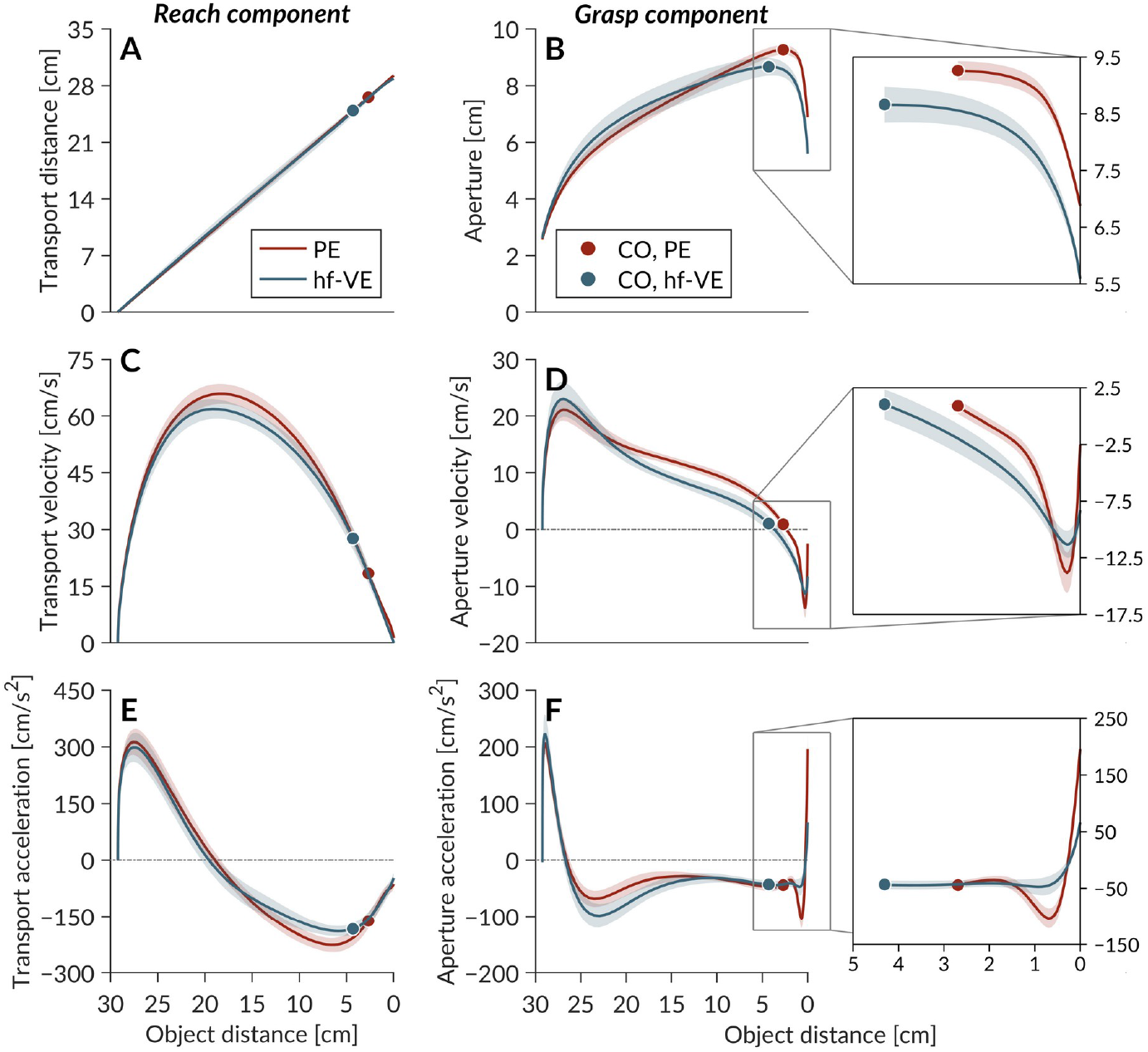
Mean spatial profiles of the reach (left) and grasp (right) components of the reach-to-grasp movements performed in PE and hf-VE, across trials for a representative object sizedistance pair (medium object, middle distance) for a representative participant. (**A**) Transport distance. (**B**) Aperture. (**C**) Transport velocity. (**D**) Aperture velocity. (**E**) Transport acceleration. (**F**) Aperture acceleration. Solid circles in each plot denote closure onset (CO). Shaded areas indicate ±1 *SEM* (*n* = 13).

### Data processing and feature extraction

All data were analyzed offline using custom MATLAB routines (Mathworks Inc., Natick, MA). For each trial, the time series data for the *x-* and *y*- dimensions were cropped from movement onset (and the moment the switch was released) to movement offset (the moment aperture velocity became zero in PE, and the moment the collision detection criterion was met in hf-VE) and filtered (6 Hz, fourth-order low-pass Butterworth filter).

For each condition (environment, object size, and object distance), mean values were calculated on a per-participant basis for the following outcome variables: (a) Movement time, defined as the interval between movement onset and movement offset; (b) Maximum aperture, which marked the initiation of closure or closure onset (CO) and which we refer to as aperture at CO; (c) Opening time, defined as the interval from movement onset to CO; (d) Closure time, defined as the interval from CO to movement offset; (e) Transport distance, defined as the distance between the initial and final positions of the wrist; (f) Opening distance, distance between the wrist’s position at movement onset and the wrist’s position at CO; (g) Closure distance, defined as the distance between the wrist’s position at CO and the object; (h) Maximum closure velocity; (i) Maximum closure deceleration; (j) Transport velocity at CO; and (k) Transport acceleration at CO.

### Statistical analyses

All analysis was performed at the level of participant means across trials, unless stated otherwise. 2 × 3 × 3 repeated measures analyses of variance (rm-ANOVAs) with within-subject factors of the Environment (PE, hf-VE), Object size (Small, Medium, Large), and Object distance (levels: Near, Middle, Far) were used to evaluate the effects of the Environment, Object size, and Object distance on each kinematic variable. All data mostly fulfilled the conditions of normality, homogeneity of variance, and sphericity; when the assumption of sphericity was not met, Greenhouse-Geisser correction was applied, and in these cases, the corrected *p*-values and the Greenhouse-Geisser correction factor (*ε*) are reported. Each test was performed in SPSS 21 (IBM Inc., Chicago, IL), and each test statistic was considered significant at the two-tailed *α* level of 0.05. All effect sizes are reported as partial eta-squared (*η*_p_^2^).

Pearson’s correlation tests were used to examine the relationship between opening/closure time and movement time in both PE and hf-VE, as well as between opening/closure distance and transport distance in both PE and hf-VE. Pearson’s correlation tests were performed over group means across participants, as well over participant means across trials, in both PE and hf-VE, to test: (1) relationship between closure distance and each state parameter; and (2) pairwise relationships among the three state parameters. Each test was performed in SPSS 21. To avoid the risk of type-I error due to multiple correlations, each test statistic was considered significant at the two-tailed *α_corrected_* level of 0.017 after correcting for multiple correlations: *α_corrected_*=1-(1–*α*)^1/*k*^, where *k* = 3 (Curtin and Schulz 1998).

To investigate which state estimates best explained variation in closure distance in PE and hf-VE, we followed an information-theoretic approach to model selection using RStudio. This approach uses the Akaike Information Criterion (AIC; or quasi-AIC (QAICc) for overdispersed data) to choose a set of plausible models from a given set of *a priori* candidate models (Burnham and Anderson 2002). AICc for a given model is computed as: 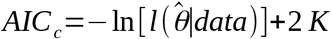, where 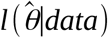 is the likelihood of the parameters 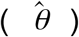 in the model and *K* = number of estimable parameters in the model. For example, a simple linear regression model, y = *a* + *bx,* has three estimable parameters: in intercept *a*, the slope b, and the variance (*s*^2^). This definition of AIC is applicable for large samples (*n*). In the smallsample case (i.e., *n/K* < 40), 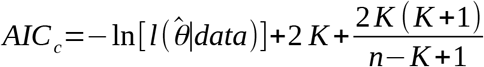. To compute QAICc, the number of model parameters is increased by 1 to account for overdispersion.

QAICc serves as an estimator of out-of-sample prediction error and thereby the relative quality of statistical models for a given set of data. QAICc estimates the quality of each model relative to each of the other models. Specifically, a smaller QAICc value reflects better performance/complexity trade-off. Thus, QAICc provides a means for selecting the model with the best performance/complexity trade-off. In the present study, we considered eight candidate models, including the null model (containing only the intercept) and all the subset models of the following full model: CD = *a* + *b*(A_CO_) + *c*(TV_CO_) + *d*(A_CO_;), by allowing the intercept to differ by participant, where CD = closure distance, A_CO_ = aperture at CO, TV_CO_ = transport velocity at CO, and TA_CO_ = transport acceleration at CO.

## Results

Figure 2 provides spatial trajectories of the reach (left) and grasp (right) components of the reach-to-grasp movements performed in PE and hf-VE for a representative object size-distance pair (medium object, middle distance). Table 1 summarizes the mean (±1*SD*) values of each movement variable across all participants (*n* = 13). Table 2 summarizes the outcomes of rm-ANOVAs. Test statistics only for significant effects are presented below (see Table 2 for non-significant outcomes.

**Table 1.** Mean (±1*SD*) values of each movement variable across all participants (*n* = 13) for reach-to-grasp movements performed in PE and hf-VE

|  |  | PE |  |  | hf-VE |  |  |
| --- | --- | --- | --- | --- | --- | --- | --- |
| Variable | Object size /distance | Near | Middle | Far | Near | Middle | Far |
| Closure time [ms] |  |  |  |  |  |  |  |
| | Small | 344±64 | 361±96 | 392±108 | 418±120 | 411±114 | 421±119 |
| | Medium | 314±63 | 324±56 | 346±78 | 413±91 | 415±111 | 441±98 |
| | Large | 300±59 | 319±58 | 314±48 | 416±128 | 423±119 | 430±97 |
| Closure distance [cm] |  |  |  |  |  |  |  |
| | Small | 3.1±1.2 | 3.1±1.0 | 3.4±1.4 | 4.7±2.0 | 5.1±2.4 | 5.6±2.7 |
| | Medium | 2.4±0.9 | 2.6±1.0 | 3.0±1.2 | 4.0±1.7 | 4.3±2.3 | 4.8±2.9 |
| | Large | 2.5±0.8 | 2.6±0.9 | 2.6±0.8 | 3.3±1.7 | 3.6±1.9 | 4.2±2.2 |
| Maximum closure velocity [cm/s] |  |  |  |  |  |  |  |
| | Small | -15.8±5.7 | -16.0±6.8 | -15.6±6.3 | -12.8±4.6 | -16.0±4.6 | -15.6±4.1 |
| | Medium | -15.8±5.5 | -15.2±6.6 | -15.3±5.6 | -13.0±4.4 | -13.9±5.4 | -13.5±3.8 |
| | Large | -14.1±4.5 | -13.7±5.0 | -13.2±4.6 | -13.2±5.9 | -12.4±5.1 | -13.0±4.4 |
| Maximum closure deceleration [cm/s<sup>2</sup>] |  |  |  |  |  |  |  |
| | Small | -143.3±63.5 | -134.7±77.9 | -134.2±62.8 | -101.7±43.7 | -105.7±36.8 | -102.0±39.6 |
| | Medium | -147.7±61.0 | -145.3±63.1 | -135.4±59.9 | -105.0±46.0 | -110.8±65.6 | -113.3±42.2 |
| | Large | -133.8±35.9 | -113.8±47.1 | -119.7±44.0 | -108.4±52.8 | -96.9±45.9 | -95.4±39.6 |

**Table 2.** Outcomes of rm-ANOVAs examining the effects of the Environment (PE, hf-VE), Object size (Small, Medium, Large), and Object distance (Near, Middle, Far) on each movement variable of reach-to-grasp movements

| Variable | Effect | *F* | df | $\epsilon^*$ | $p^\dagger$ | $\eta_p^2$ |
| --- | --- | --- | --- | --- | --- | --- |
| Closure time [ms] |  |  |  |  |  |  |
|  | Environment | 19.24 | 1,12 |  | **0.001** | 0.62 |
|  | Object size | 3.77 | 2,12 |  | **0.038** | 0.24 |
|  | Object distance | 11.29 | 2,12 |  | **< 0.001** | 0.49 |
|  | Environment × Object size | 7.69 | 2,12 |  | **0.003** | 0.39 |
|  | Environment × Object distance | 1.11 | 2,12 | 0.69 | 0.329 | 0.09 |
|  | Object size × Object distance | 0.78 | 4,12 | 0.53 | 0.475 | 0.06 |
|  | Environment × Object size × Object distance | 0.76 | 4,12 |  | 0.559 | 0.06 |
| Closure distance [cm] |  |  |  |  |  |  |
|  | Environment | 15.63 | 1,12 |  | **0.002** | 0.57 |
|  | Object size | 20.25 | 2,12 |  | **0.001** | 0.63 |
|  | Object distance | 13.66 | 2,12 | 0.70 | **0.007** | 0.53 |
|  | Environment × Object size | 6.77 | 2,12 |  | **0.005** | 0.36 |
|  | Environment × Object distance | 2.64 | 2,12 |  | 0.092 | 0.18 |
|  | Object size × Object distance | 0.31 | 4,12 | 0.65 | 0.788 | 0.03 |
|  | Environment × Object size × Object distance | 0.66 | 4,12 |  | 0.622 | 0.05 |
| Maximum closure velocity [cm/s] |  |  |  |  |  |  |
|  | Environment | 1.77 | 1,12 |  | 0.208 | 0.13 |
|  | Object size | 5.84 | 2,12 |  | **0.009** | 0.33 |
|  | Object distance | 0.12 | 2,12 |  | 0.886 | 0.01 |
|  | Environment × Object size | 0.62 | 2,12 | 0.69 | 0.492 | 0.05 |
|  | Environment × Object distance | 1.83 | 2,12 |  | 0.182 | 0.13 |
|  | Object size × Object distance | 1.16 | 2,12 |  | 0.343 | 0.09 |
|  | Environment × Object size × Object distance | 0.80 | 4,12 |  | 0.533 | 0.06 |
| Maximum closure deceleration [cm/s<sup>2</sup>] |  |  |  |  |  |  |
|  | Environment | 8.55 | 1,12 |  | **0.015** | 0.42 |
|  | Object size | 3.52 | 2,12 |  | **0.046** | 0.23 |
|  | Object distance | 2.86 | 2,12 |  | 0.077 | 0.19 |
|  | Environment × Object size | 0.76 | 2,12 | 0.65 | 0.428 | 0.06 |
|  | Environment × Object distance | 1.78 | 2,12 |  | 0.179 | 0.13 |
|  | Object size × Object distance | 1.67 | 2,12 |  | 0.173 | 0.12 |
|  | Environment × Object size × Object distance | 0.14 | 4,12 |  | 0.965 | 0.01 |

\*Greenhouse-Geisser correction factor for violation of the assumption of sphericity.
†Boldfaced values indicate statistical significance at the two-tailed alpha level of 0.05.

### Closure was slower to occur, lasted longer and was initiated farther from the object in hf-VE than in PE

Closure lasted longer in hf-VE than in PE by mean±1 *SEM* = 86±19 ms (rm-ANOVA: *F*_1,12_ = 19.24, *p* = 0.001, *η*_p_^2^ = 0.62; Fig. 3A, left). Closure depended on object size (*F*_2,12_ = 3.77, *p* = 0.038, *η*_p_^2^ = 0.24), and this effect was mediated by the environment (*F*_2,12_ = 7.69, *p* = 0.003, *η*_p_^2^ = 0.39; Fig. 3A, left). Bonferroni’s post-hoc tests revealed that closure time reduced with object size in PE (mean±1 *SEM* difference: small vs. large object: 54±14 ms, *p* = 0.006) but closure time did not depend on object size in hf-VE (*p* > 0.05, Fig. 3B, left). Closure time increased with object distance (*F*_2,12_ = 11.29, *p* < 0.001, *η_p_*^2^ = 0.49) and this effect did not depend on the environment (*p* > 0.05; Fig. 3A, right).

**Fig. 3.**
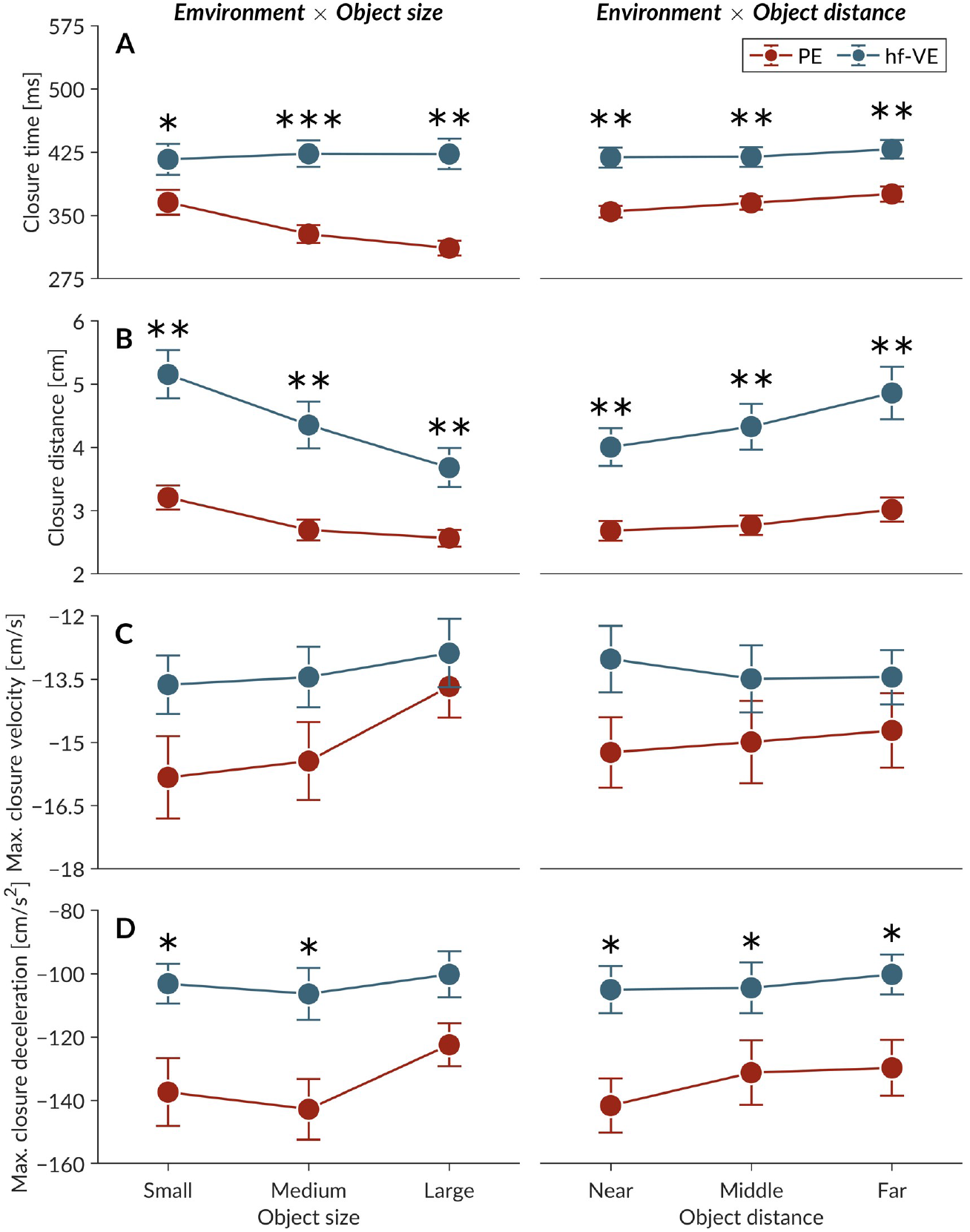
Values of closure variables for the reach-to-grasp movements performed in PE and hf-VE. (**A**) Closure time. (**B**) Closure distance. (**C**) Maximum closure velocity. (**D**) Maximum closure deceleration. Error bars indicate ±1*SEM* (*n* = 13), and individual points represent the participant means for the near, middle, and far distances (left) and the small, medium, and large sizes. *\*p* < 0.05; *\*\*p* < 0.005; \*\*\**p* < 0.001.

Closure was initiated farther from the object in hf-VE than in PE by mean±1 *SEM* = 1.6±0.4 cm (*F*_1,12_ = 15.63, *p* = 0.002, *η_p_*^2^ = 0.57; Fig. 3B). Closure distance reduced with object size (*F*_2,12_ = 20.25, *p* = 0.001, *η_p_*^2^ = 0.63), and this effect was mediated by the environment (*F*_2,12_ = 6.77, *p* = 0.005, *η_p_*^2^ = 0.36; Fig. 3B, left). Bonferroni’s post-hoc tests revealed that this reduction in closure distance with object size was stronger in hf-VE (mean±1 *SEM* difference: small vs. large object: 1.5±0.3 cm, *p* < 0.001; medium vs. large object: 0.8±0.3 cm, *p* = 0.028) than in PE (mean±1 *SEM* difference: small vs. large object: 0.7±0.7 cm, *p* = 0.005; medium vs. large object: 0.5±0.1 cm, *p* = 0.009, Fig. 3B, left). Closure distance increased with object distance (*F*_2,12_ = 13.66, *p* = 0.007, *η_p_*^2^ = 0.53) and this effect did not depend on the environment (*p* > 0.05; Fig. 3B, right).

Maximum closure velocity increased with object size (*F*_2,12_ = 5.84, *p* = 0.009, *η_p_*^2^ = 0.33) but did not differ between the two environments (*p* > 0.05; Fig. 3C, left). Maximum closure deceleration increased with object size (*F*_1,12_ = 3.52, *p* = 0.046, *η_p_*^2^ = 0.23), as well as it was lower in hf-VE than in PE by mean±1 *SEM* = 31.0±10.6 cm/s^2^ (*F*_1,12_ = 8.55, *p* = 0.015, *η_p_*^2^ = 0.42; Fig. 3D, left).

Performing reach-to-grasp movements in hf-VE weakened the dependence of closure time on object size and strengthened the dependence of closure distance on object size, with an amplifying effect of hf-VE on both closure time and closure distance. Critically, closure time and closure distance showed the same relationship with object distance observed by Rand et al. (2006b), again, with an amplifying effect of hf-VE. In summary, closure was slower to occur, lasted longer and was initiated farther from the object in hf-VE than in PE.

### Closure distance did not scale with transport distance in either PE or hf-VE

Previously it had been suggested that the scaling of closure time with transport time but not closure distance with transport distance indicates that initiation of aperture closure is based primarily on spatial rather than temporal parameters of the movement (Rand et al. 2006a, b). To examine the stability of closure time and distance in our data, we tested the association between opening/closing time and total movement time, and between opening/closing distance and total transport distance. Opening time showed a strong positive relationship with movement time in both PE (Pearson’s correlation, *ρ* = 0.85, *p* < 0.001; Fig. 4A) and hf-VE (*ρ* = 0.76, *p* < 0.001; Fig. 4B), and closing time showed a weak positive relationship with movement time in both PE (*ρ* = 0.50, *ρ* < 0.001; Fig. 4A) and hf-VE (*ρ* = 0.43, *p* < 0.001; Fig. 4B), indicating opening and closing time increased with increased movement time. The slopes of the regression lines for both opening time (0.72 in PE vs. 0.71 in hf-VE) and closing time (0.27 in PE vs. 0.29 in hf-VE) were similar between the two environments. Opening distance showed a strong positive relationship with transport distance in both PE (*ρ* = 0.98, *p* < 0.001; Fig. 4C) and hf-VE (*ρ* = 0.91, *p* < 0.001; Fig. 4D). In contrast, closing distance did not show a significant relationship with transport distance in PE (*ρ* = 0.11, *p* = 0.505; Fig. 4C) or hf-VE (*ρ* = 0.14, *p* = 0.306; Fig. 4D). The slopes of the regression lines for both opening distance (0.98 in PE vs. 0.94 in hf-VE) and closing distance (0.024 in PE vs. 0.060 in hf-VE) were similar between the two environments. The relative stability of closure distance compared to closure time with varying total movement characteristics confirms that VE and hf-PE show a similar spatiotemporal relationships between opening and closing and that the relationship observed in both enviornments is similar to that observed by Rand et al.’s (2006a, b).

**Fig. 4.**
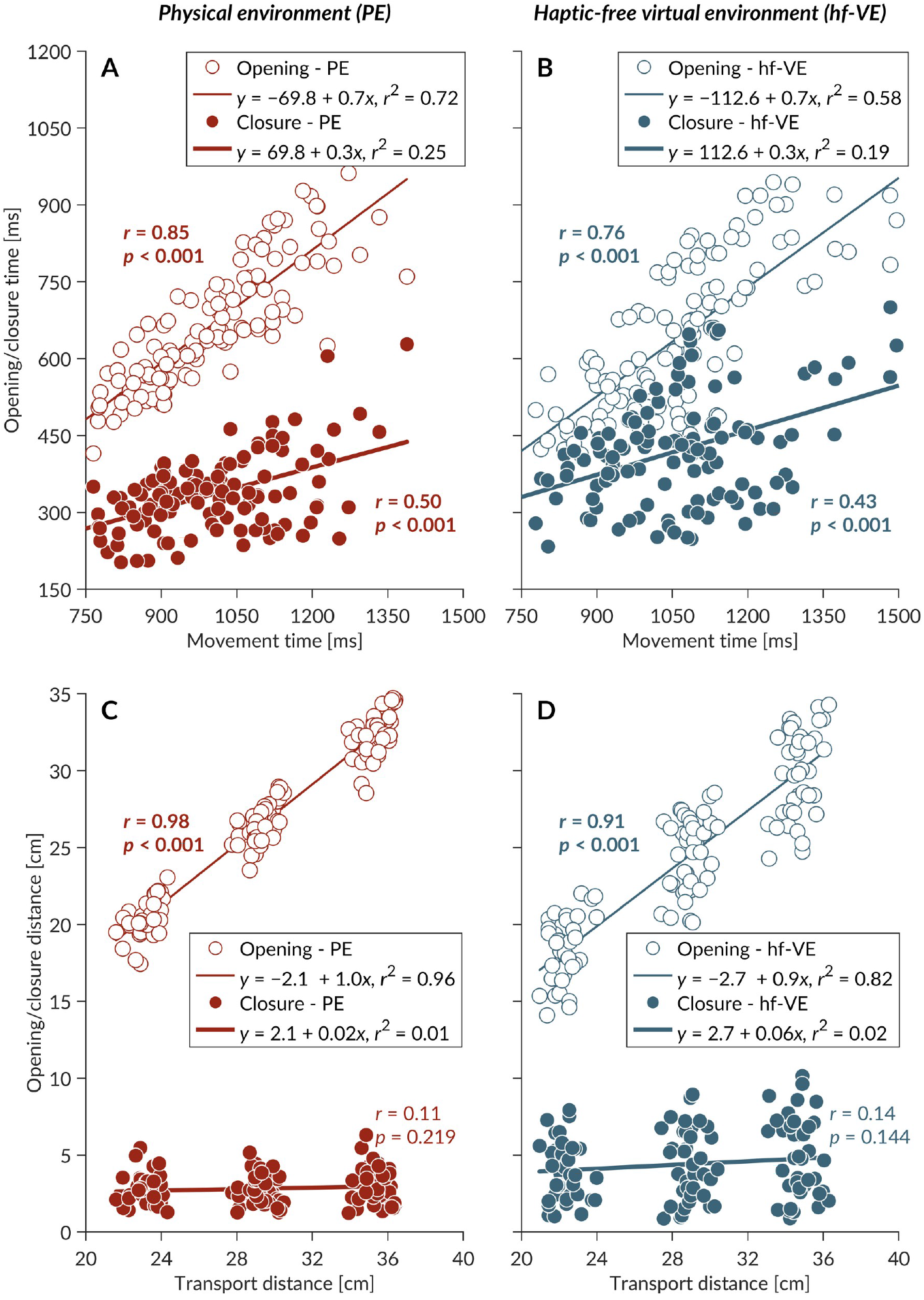
The relationship between opening/closure time and movement time, and between opening/closure distance and transport distance, across all object sizes and distances in PE and hf-VE. (**A**) Opening/closure time vs. movement time in PE. (**B**) Opening/closure time vs. movement time in hf-VE. (**C**) Opening/closure distance vs. transport distance in PE. (**D**) Opening/closure distance vs. transport distance in hf-VE.

### Similar control laws governed the initiation of closure in PE and hf-VE

Of the eight models used to identify the control law governing the initiation of closure in PE, only two models yielded nonzero weights, *W*_i_, (the probability that a given model is the best model among all models). Support for the first model was 999 times stronger than the second model (evidence ratio = *W*_1_/*W*_2_ = 0.9900/0.0010 = 999; *χ^2^* = 16.66, *p* < 0.001, Table 3). The first model yielded both the best fit (log-likelihood closest to zero) and the best performance-to-complexity tradeoff (lowest QAICc). This model included a coefficient for A_CO_, TV_CO_, and TA_CO_, supporting the idea presented by Rand and coworkers (Rand et al. 2006a, b) that the distance at closure is predicted by a linear function combining the state estimates of transport velocity, transport acceleration, and aperture.

**Table 3.** Summary of model selection performed on participant means across trials.

| Physical environment (PE) |  |  |  |  |  |  |  |  |  |
| --- | --- | --- | --- | --- | --- | --- | --- | --- | --- |
| Model (i) | $B$ | $A_{CO}$ | $TV_{CO}$ | $TA_{CO}$ | $K$ | Log-Likelihood | QAICc | $\Delta_i$ | $w_i$ |
| 8 | 1.46 | −0.084 | 0.13 | 0.0030 | 6 | −25.56 | 63.9 | 0.00 | 0.9990 |
| 6 | 1.37 | −0.092 | 0.12 |  | 5 | −33.89 | 78.3 | 14.43 | 0.0010 |
| 7 | 0.55 |  | 0.14 | 0.0034 | 5 | −34.95 | 80.4 | 16.56 | 0.00 |
| 5 | 0.34 |  | 0.12 |  | 4 | −43.91 | 96.2 | 32.30 | 0.00 |
| 4 | 3.81 | −0.18 |  | −0.0042 | 5 | −104.88 | 220.3 | 156.43 | 0.00 |
| 2 | 4.60 | −0.19 |  |  | 4 | −110.97 | 230.3 | 166.42 | 0.00 |
| 3 | 2.03 |  |  | −0.0048 | 4 | −116.60 | 241.6 | 177.67 | 0.00 |
| 1 | 2.82 |  |  |  | 3 | −123.00 | 252.2 | 188.33 | 0.00 |
| MAP | 1.45 | −0.084 | 0.13 | 0.0030 |  |  |  |  |  |
| 2.5% CI | 0.94 | −0.12 | 0.12 | 0.0016 |  |  |  |  |  |
| 97.5% CI | 1.96 | −0.047 | 0.15 | 0.0044 |  |  |  |  |  |
| Haptic-free virtual environment (hf-VE) |  |  |  |  |  |  |  |  |  |
| Model (i) | $B$ | $A_{CO}$ | $TV_{CO}$ | $TA_{CO}$ | $K$ | Log-Likelihood | QAICc | $\Delta_i$ | $w_i$ |
| 7 | −0.42 |  | 0.19 | 0.0048 | 5 | −85.05 | 124.0 | 0.00 | 0.73 |
| 8 | −0.74 | 0.028 | 0.20 | 0.0050 | 6 | −84.86 | 126.0 | 2.01 | 0.27 |
| 5 | −0.74 |  | 0.18 |  | 4 | −94.61 | 134.3 | 10.29 | 0.0040 |
| 6 | −0.43 | −0.027 | 0.17 |  | 5 | −94.45 | 136.1 | 12.30 | 0.0020 |
| 4 | 7.20 | −0.47 |  | −0.0080 | 5 | −180.95 | 249.5 | 125.44 | 0.00 |
| 2 | 8.77 | −0.49 |  |  | 4 | −187.73 | 560.1 | 132.09 | 0.00 |
| 3 | 2.85 |  |  | −0.0091 | 4 | −195.24 | 265.9 | 141.92 | 0.00 |
| 1 | 4.40 |  |  |  | 3 | −202.08 | 272.7 | 148.68 | 0.00 |
| MAP | −0.50 | 0.0073 | 0.19 | 0.0048 |  |  |  |  |  |
| 2.5% CI | −1.31 | −0.056 | 0.18 | 0.0028 |  |  |  |  |  |
| 97.5% CI | 0.30 | 0.11 | 0.21 | 0.0070 |  |  |  |  |  |

Parameters include the intercept ( $B$ ); the coefficients for each of the continuous predictors: aperture at CO ( $A_{CO}$ ), transport velocity at CO ( $TV_{CO}$ ), and transport acceleration at CO ( $TA_{CO}$ ); the number of free parameters ( $K$ ); Log-Likelihood; the Akaike information criterion corrected for overdispersion (QAICc); the difference in QAICc between the $i$ th model and the best model ( $\Delta_i$ ); and model weight or the probability that a given model is the best model among all models ( $w_i$ ). Models are arranged in order from best (lowest $\Delta_i$ ) to worst (highest $\Delta_i$ ). The bottom rows describe the model-averaged parameter estimates (MAP) with lower (2.5%) and upper (97.5%) bounds of 95% confidence intervals (CI).

Of the eight models used to identify the control law governing the initiation of closure in hf-VE, only two models yielded non-negligible *w*_i_. Support for the first model was 3.2 times stronger than the second model (evidence ratio = *W*_1_/*W*_2_ = 0.73/0.27 = 2.50; Table 3). While the model with the best fit (model 2, log-likelihood closest to zero) included a coefficient for all three variables, the model with the best performance-to-complexity tradeoff (model 1, lowest QAICc) did not include a coefficient for aperture at CO. Nonetheless, the first and second models yielded close goodness of fit (log-likelihood values: −85.05 and −84.86, respectively) and the inclusion of A_CO_ in the model did not significantly improve the fit (*χ*^2^ = 0.38, *p* = 0.537). In short, the initiation of closure in hf-VE was not as dependent on aperture at CO as it did in PE.

To further explore the contribution of different state parameters to the respective control laws, we analyzed the relationship between the values of closure distance and each state parameter averaged across all participants. Closure distance showed a negative relationship with aperture at CO in both environments (PE: Pearson’s *r* = −0.78, *p* = 0.013; hf-VE: *r* = −0.81, *p* = 0.008; Fig. 5A, left), a strong positive relationship with transport velocity at CO (PE: *r* = 0.94, *p* < 0.001; hf-VE: *r* = 0.98, *p* < 0.001; Fig. 5B, left), and a weak negative relationship with transport acceleration at CO (PE: *r* = −0.60, *p* = 0.088; hf-VE: *r* = −0.64, *p* = 0.067; Fig. 5C, left). The strong relationship between closure distance and transport velocity at CO in both PE and hf-VE indicates that it is the most important state parameter contributing to the control laws. This result confirms that of the model selection in that the model with the best fit and performance-to-complexity tradeoff for PE, and all four models with nonzero weight, *W*_i_, for hf-VE, included a coefficient for transport velocity at CO (Table 3).

**Fig. 5.**
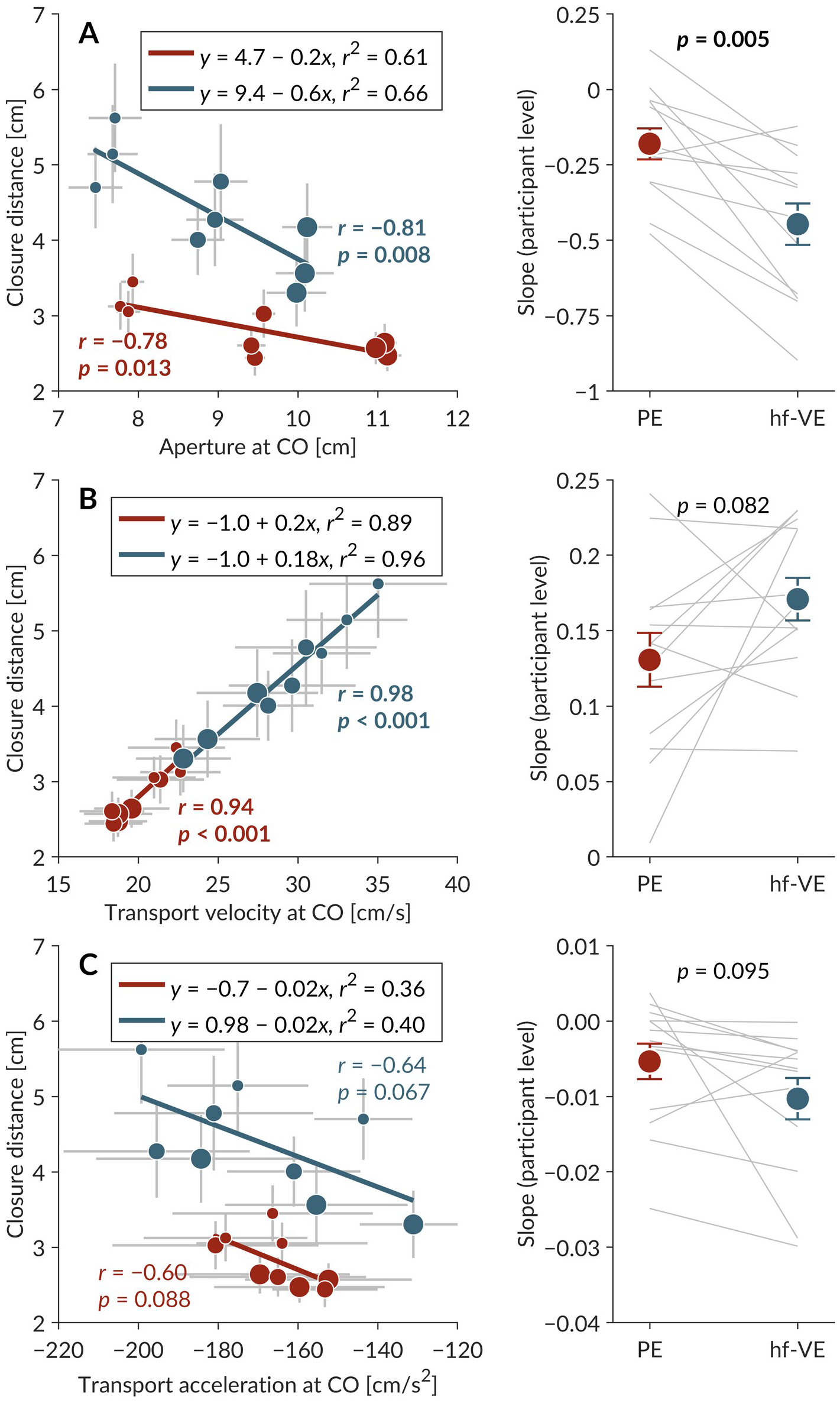
Relationship between closure distance and each state parameter in across all object sizes and distances PE (red) and hf-VE (blue) at the level of group means across participants. (**A**) Closure distance and aperture at CO. (**B**) Closure distance and transport velocity at CO. (**C**) Closure distance and transport acceleration at CO. Left panels show pairwise relationship in PE and hf-VE. Right panels show comparisons of the participant-level slope of the regression line between PE and hf-VE using paired samples *t*-test. To facilitate statistical comparison of slopes between the two environments, the comparisons of participant-level slopes confirm that differences in the respective pairwise relationships between PE and hf-VE observed at the group level also held at the participant level. Small, medium, and large circles indicate group means across participants for the small, medium, and large objects, respectively, placed at each of the three distances. Horizontal and vertical bars indicate ±1 *SEM* (*n* = 13).

To substantiate that the state estimate of aperture contributed to the variability in the control law governing the initiation of closure between PE and hf-VE, we compared the relationship between closure distance and each state parameter between PE and hf-VE at the level of group means across participants. In both PE and hf-VE, closure distance showed a strong negative relationship with aperture at CO (PE: Pearson’s *r* = −0.78, *p* = 0.013; hf-VE: *r* = −0.81, *p* = 0.008; Fig. 5A, left) and a very strong positive relationship with transport velocity at CO (PE: *r* = 0.94, *p* < 0.001; hf-VE: *r* = 0.98, *p* < 0.029; Fig. 5B, left) but did not show a significant relationship with transport acceleration at CO (PE: *r* = −0.060, *p* = 0.088; hf-VE: *r* = −0.064, *p* = 0.067; Fig. 5C, left). At the level of participant means across trial, the slope of the regression line between closure distance and aperture at CO was steeper in hf-VE than in PE (paired *t*-test: *t*_2,13_ = 4.40, *p* = 0.029; Fig. 5A, right). Critically, the slope of the regression line did not differ for transport velocity at CO (*t*_2,13_ = −1.90, *p* = 0.082; Fig. 5B, right) and transport acceleration at CO (*t*_2,13_ = 1.81, *p* = 0.095; Fig. 5C, right). The comparisons of participant-level slopes confirm that differences in the respective pairwise relationships between PE and hf-VE observed at the group level also held at the participant level.

Finally, to test whether a difference in the relationship between aperture at CO and transport velocity at CO might be a possible reason for why the initiation of closure in hf-VE did not depend on aperture at CO as it did in PE, we compared the pairwise relationships among the three state parameters between PE and hf-VE at the level of group means across participants. In both PE and hf-VE, aperture at CO showed a strong negative relationship with transport velocity at CO (PE: Pearson’s *r* = −0.72, *p* = 0.023; hf-VE: *r* = −0.88, *p* = 0.002; Fig. 6A, left), although the relationship in PE was not significant after correcting for multiple correlations. Aperture at CO did not show a relationship with transport acceleration at CO (PE: *r* = 0.38, *p* = 0.309; hf-VE: *r* = 0.21, *p* = 0.594; Fig. 6B, left). After correcting for multiple correlations, transport velocity at CO did not show a relationship with transport acceleration at CO in both PE and hf-VE (PE: *r* = −0.72, *p* = 0.029; hf-VE: *r* = −0.62, *p* = 0.072; Fig. 6C, left). At the level of participant means across trials, the slope of the regression line between aperture at CO and transport velocity at CO was steeper in hf-VE than in PE (paired *t*-test: *t*_2,13_ = 4.47, *p* < 0.001; Fig. 6A, right). Critically, the slope of the regression line between aperture at CO and transport acceleration at CO did not differ between PE and hf-VE (*t*_2,13_ = −0.11, *p* = 0.917; Fig. 6B, right), and the slope of the regression line between transport velocity at CO and transport acceleration at CO was only marginally steeper in hf-VE than in PE (*t*_2,13_ = 2.21, *p* = 0.047; Fig. 6C, right).

**Fig. 6.**
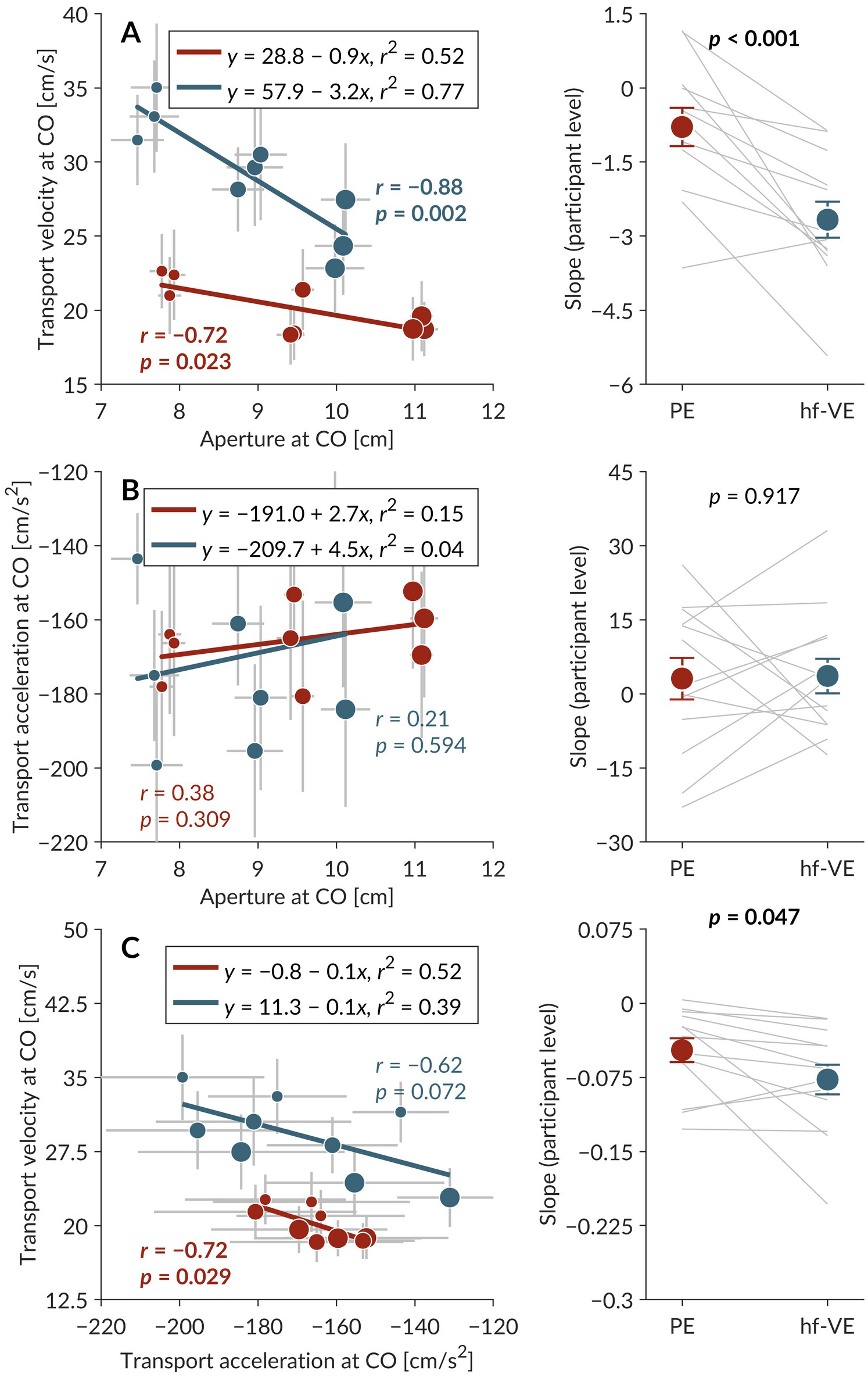
Pairwise relationships among the three state parameters across all object sizes and distances in PE (red) and hf-VE (blue) at the level of group means across participants. (**A**) Transport velocity at CO and aperture at CO. (**B**) Transport acceleration at CO and aperture at CO. (**C**) Transport velocity at CO and transport acceleration at CO. Left panels show pairwise relationship in PE and hf-VE. To facilitate statistical comparison of slopes between the two environments, right panels show comparisons of the participant-level slope of the regression line between PE and hf-VE using paired samples *t*-test. The comparisons of participant-level slopes confirm that differences in the respective pairwise relationships between PE and hf-VE observed at the group level also held at the participant level. Small, medium, and large circles indicate group means across participants for the small, medium, and large objects, respectively, placed at each of the three distances. Horizontal and vertical bars indicate ±1*SEM* (*n* = 13).

## Discussion

In the present study, reanalysis of the Furmanek et al. (2019) data indicated that the closure phase was also initiated farther from the object, with lower maximum closure deceleration, in immersive hf-VE than in PE. These differences presumably reflect the fact that when reaching to grasp in hf-VE, the voluntary effort needed to actively bring the fingers to a stop at the interface of a virtual object. PE does not impose this physical constraint, as the fingers automatically stop when they contact the physical object. However, the goal of this study was not merely to identify differences in the initiation of closure between the two environments; the goal of the present study was to investigate potential differences and similarities in the control laws governing the initiation of closure in PE and an immersive hf-VE. To do so, a linear regression model—previously suggested by Rand et al. (2008) to predict the distance from the object at which the closure is initiated from the object from the limb state parameters of transport velocity, transport acceleration, and maximum aperture— was applied to kinematic data from each environment. The model comparisons yielded two important insights: (1) Transport velocity was the dominant predictor of closure distance in both environments, and the relationship between transport velocity at CO and closure distance was nearly identical between environments, and (2) Aperture was less informative to the control law in hf-VE.

In hf-VE, two factors significantly influence the speed-accuracy tradeoff in reach-to-grasp movements. The first is the well-documented finding that VR increases the perceptual uncertainty about object distance and size (Armbrüster et al. 2008; Ogawa et al. 2018; Harris et al. 2019). The increase in perceptual uncertainty about the spatial properties of objects, which is hypothesized to cause slower movements in hf-VE (Viau et al. 2004; Magdalon et al. 2011), was likely a major factor in smaller maximum transport velocity found in the present study. The second factor influencing the speed-accuracy tradeoff is the precision demand of grasping. The lack of physical properties of the object in hf-VE not only requires the voluntary effort needed to actively bring the fingers to a stop at the interface of a virtual object, but also increases the demand for precision of the finger aperture in the closure phase of the movement. Higher precision demands for grasping the smaller object drove a negative relationship between closure distance and aperture at CO (Table 1; Fig 5A, left), and scaling of closure distance to the aperture at CO was significantly greater in hf-VE than in PE (Fig 5A right). This result agrees with a previous study that indicated aperture closure distance systematically increased task difficulty when the availability of vision modified task demands during prehension (Rand et al. 2007).

Investigations of the speed-accuracy tradeoff for reach to grasp have shown that the full reach-to-grasp movement is poorly described by the Fitts law (Bootsma et al. 1994). Instead, it may be better characterized by Welford’s (1968) theoretical position that object distance and size place separate constraints upon movement time (McIntosh et al. 2018). However, Rand et al. (2008) have suggested that the precision of the closure phase specifically may be reasonably well described by principles of speed-accuracy tradeoff as described by the standard form of the Fitts’ law (Fitts 1954). Based on the finding that controlled manipulations of movement speed did not alter the relationship between transport velocity at CO and closure distance, Rand et al. (2006b) reasoned that variability in the opening process might be more acceptable as long as the transport-aperture coordination at the time of the phase transition to closure is tightly controlled. Our data support this assertion by Rand for movements conducted in hf-VE with one notable exception. Rand et al. (2006b) found faster movements (i.e., movements with greater maximum transport velocity) were associated with higher transport velocity at CO, and the closure initiated farther from the object. In contrast, we observed that the movements performed in hf-VE, despite having significantly lower maximum transport velocity than those performed in PE (Furmanek et al. 2019), were characterized by higher transport velocity at CO and closure distance farther from the object. Indeed, this finding appears to be consistent with previous empirical results suggesting that precision demands of grasping are associated with an extended deceleration phase, suggesting an important role for feedback-based correction (MacKenzie et al. 1987; Bootsma et al. 1994; MacKenzie and Graham 1997). We suggest that a parsimonious explanation for this difference between our result and that of Rand et al. (2006b) is that movement plan at the time of movement initiation factored the specific task and precision demands of the closure in hf-VE to extend the relationship between transport velocity at CO and closure distance learned over a lifetime of practice in PE. The participants treated the virtual object as if it were a physical object with greater precision demands for grasping (Gentilucci et al. 1991; Iyengar et al. 2009; Magdalon et al. 2011). The striking similarity of the relationship between transport velocity at CO and closure distance offers valuable insights into the critical importance of preserving this aspect of the phase transition between the opening and closing of the grasp for successful reach to grasp. We suggest that the preservation of this relationship may be a critical benchmark for any VE in which reach-to-grasp tasks are performed.

Our second insight was that aperture was less informative to the control law in hf-VE. This, too, appears to relate directly to uncertainty about spatial properties of the object and precision demands of the task. It has been shown that calibration of grasp size with reach distance allows executing naturalistic reach-to-grasp movements even without haptic terminal feedback (e.g., Bingham et al. 2007). A lack of any opportunity to calibrate grasp size with reach distance in the present study might have led to uncertainty about object size in hf-VE. Indeed, the finding that closure distance scaled differently with object size in PE and hf-VE speaks directly to the perceptual uncertainty about object size in hf-VE. Furthermore, aperture at CO was less independent in hf-VE than in PE, ultimately resulting in the aperture at CO having a weaker influence on the initiation of closure. Indeed, the strong negative relationship between transport velocity and aperture at CO in hf-VE (Fig. 6A, left) speaks to the deliberate slowing of hand transport in situations with higher precision demand, as expected of hf-VE.

In the original study (Furmanek et al. 2019), we reported that reach-to-grasp movements were similarly coordinated in both PE and hf-VE, except that the closure lasted longer hf-VE than in PE. The present reanalysis builds on this finding and shows that despite lasting longer in hf-VE than in PE, closure initiation is based on a control law that is common to both environments. Given that this control law directly speaks to the coordination between the reach and grasp components, the present findings suggest that at least this critical aspect of naturalistic reach-grasp coordination is preserved in our immersive hf-VE even in the absence of terminal haptic feedback.

These findings are especially relevant to studying the role that terminal feedback in naturalistic reach-to-grasp movements and to reproducing naturalistic reach-to-grasp movements in immersive hf-VEs. When reaching to grasp a physical object, one of the essential roles of terminal haptic feedback is to convey, via somatosensory signals, that the object has been grasped successfully. This feedback may benefit subsequent actions by updating and recalibrating forward models controlling reach-to-grasp movements to grasp size and reach distance can be scaled appropriately (Coats et al. 2008; Cavina-Pratesi and Hesse 2013). Notably, in the present study, when a collision detection algorithm detected that both 3D spheres (denoting the 3D position of the fingertips in hf-VE) had come into contact with the virtual object, the object’s color changed from blue to red to provide [terminal] visual feedback that the object had been grasped. This visual feedback could presumably substitute for the haptic feedback available upon grasping a physical object, and thus the reach-to-grasp movements were coordinated similarly in both PE and hf-VE (Prachyabrued and Borst 2014; Geiger et al. 2018). It should be noted that only visual feedback of object contact was tested and that it has been reported that auditory feedback leads to faster movements in VE than visual feedback (Zahariev and MacKenzie 2007; Zahariev and Mackenzie 2008), so we cannot extrapolate our results to audio feedback of hf-VE grasp. Future work may investigate how bio-inspired collision detection algorithms and a combination of multisensory feedback of object contact can help to bridge the remaining gap between prehensile actions performed in PE and immersive hf-VE.

To conclude, the present findings addresses the concern that the planning, execution, and control of manual actions in hf-VEs than in PE may involve distinct mechanisms (Harris et al. 2019). Such a distinction implies that VR might have certain limitations in its utility for training and rehabilitation. However, we show that albeit with different weights in hf-VE, a control law that involves sensorimotor integration of the internal state estimates of the velocity and acceleration of the arm and perceptual estimate object size dictates the initiation of closure in both PE and hf-VE. Together, the present findings suggest that hf-VEs can support a critical aspect of naturalistic coordination of reach-to-grasp movements: the initiation of closure, which is governed by a specific control law. Hence, haptic-free immersive virtual environments can be a viable option for low-cost home-based personalized training and rehabilitation, as well as for behavioral and neurophysiological investigations of reach-to-grasp movements and it’s underlying control mechanisms.

## Acknowledgments

This work was supported by NIH grants #R01NS085122 and #2R01HD058301, and NSF grants #CBET-1804550 and #CMMI-M3X-1935337, to Eugene Tunik. We thank Alex Hunton and Samuel Berin for developing the VR platform.

## Author contributions

M.M., M.Y., M.P.F., and G.T. conceived and designed research; M.P.F. performed experiments; M.M., M.P.F., and M.P.F. analyzed data; M.M., M.Y., M.P.F., and G.T. interpreted results of experiments; M.M. prepared figures; M.M. drafted manuscript; M.M., M.Y., M.P.F., and G.T. edited and revised manuscript; M.M., M.Y., M.P.F., and G.T. approved final version of manuscript.

## Conflict of interest

The authors declare that no competing interests exist.

## Notes

### Competing Interest Statement

The authors have declared no competing interest.

